# Farnesoid X receptor agonism prevents neutrophil extracellular traps via reduced sphingosine-1-phosphate in chronic kidney disease

**DOI:** 10.1101/2023.03.01.530697

**Authors:** Bryce A. Jones, Komuraiah Myakala, Mahilan Guha, Shania Davidson, Sharmila Adapa, Isabel Lopez Santiago, Isabel Schaffer, Yang Yue, Jeremy Allegood, Lauren A. Cowart, Xiaoxin X. Wang, Avi Z. Rosenberg, Moshe Levi

**Author notes:** Corresponding author: Moshe Levi 3900 Reservoir Rd NW Basic Science Building, Room 353 Washington, DC 20007. **Significance Statement:** Many preclinical studies have shown that the farnesoid X receptor (FXR) reduces renal inflammation, but the mechanism is poorly understood. This report identifies FXR as a novel regulator of neutrophilic inflammation and NETosis via the inhibition of sphingosine-1-phosphate signaling in male but not female mice. NETosis is also identified in human Alport kidney biopsies. A better understanding of this signaling axis may lead to novel treatments that prevent renal inflammation.

## Abstract

**Background:** Activation of the farnesoid X receptor (FXR) reduces renal inflammation, but the underlying mechanisms remain elusive. Neutrophil extracellular traps (NETs) are webs of DNA formed when neutrophils undergo specialized programmed cell death (NETosis). Sphingosine-1-phosphate (S1P) is a signaling lipid that stimulates NETosis via its receptor on neutrophils. Here, we identify FXR as a negative regulator of kidney NETosis via repressing S1P signaling in male but not female mice.

**Methods:** We determined the effects of the FXR agonist obeticholic acid (OCA) in mouse models of adenosine phosphoribosyltransferase deficiency and Alport syndrome. We assessed renal NETosis by immunofluorescence in these models and in biopsies from patients with Alport syndrome (6 male, 9 female). We also inhibited de novo sphingosine production in Alport mice to show a causal relationship between S1P signaling and renal NETosis.

**Results:** Renal FXR activity is greatly reduced in both models, and OCA prevents kidney fibrosis, inflammation, and lipotoxicity. OCA reduces renal neutrophilic inflammation and NETosis in male adenine and Alport mice, but not in female adenine mice. Extensive NETosis was also identified in human Alport kidney biopsies. Kidney sphingosine kinase 1 (Sphk1) expression is increased in mice with kidney disease and reduced by OCA in male but not female mice. Also, Sphk1 expression correlates with NETosis in male but not female mice. Short-term inhibition of sphingosine synthesis reduces neutrophilic inflammation and NETosis.

**Conclusion:** FXR agonism represses kidney Sphk1 expression in male but not female mice. This inhibits renal S1P signaling, thereby reducing neutrophilic inflammation and NETosis in a sex-dependent manner.

## Introduction

Inherited kidney diseases are a collection of over 150 disorders, most arising from single-gene mutations, that are responsible for over 10% of patients on renal replacement therapy.^1^ We investigate activation of the farnesoid X receptor (FXR) in models of adenosine phosphoribosyl transferase (APRT) deficiency and Alport syndrome, both rare causes of chronic kidney disease (CKD).

FXR is a nuclear receptor that is endogenously activated by bile acids, and obeticholic acid (OCA) is a clinically approved FXR agonist.^2–4^ Numerous preclinical studies have shown that FXR protects against kidney disease.^5–12^ Briefly, FXR activation decreases renal fibrosis, inflammation, lipotoxicity, oxidative stress, and endoplasmic reticulum stress.^13^ However, mechanistic studies on how FXR agonism reduces renal inflammation are lacking. We address this gap herein.

Neutrophils are innate immune cells, and one mechanism whereby neutrophils slow infection progression is to immobilize microbes within neutrophil extracellular traps (NETs). NETs are stringy webs of decondensed DNA and antimicrobial proteins that are released into the extracellular milieu during NETosis, a form of programmed cell death.^14, 15^ However, aberrant NETosis has been reported in sterile inflammation, including some kidney diseases.^16^ No study thus far has investigated NETs in models of APRT deficiency or Alport syndrome, nor has the role of FXR in NETosis been investigated.

Sphingosine-1-phosphate (S1P) is a pro-inflammatory signaling lipid produced from sphingosine by its two kinases: Sphk1 and Sphk2. S1P is best understood in the context of T-cells,^17–20^ but emerging research indicates that S1P also promotes neutrophilic inflammation. S1P increases neutrophil chemotaxis^21–24^ and activates NETosis via its G protein-coupled receptors on neutrophils.^25, 26^ Inhibiting S1P signaling stimulates neutrophil apoptosis and clearance by macrophages, promoting the resolution of neutrophilic inflammation.^23, 27^ Although a few initial studies indicated that S1P may be beneficial to the kidney,^28, 29^ many recent studies using Sphk1-null mice showed that inhibition of Sphk1 protects the kidney.^30–34^ Of relevance to the current study, a recent report showed that FXR repressed Sphk1 expression in the liver (supp. info. in ^35^).

Taken together, we hypothesized that FXR agonism represses kidney Sphk1 expression and lowers S1P, thereby reducing neutrophilic inflammation and NETosis in models of APRT deficiency and Alport syndrome.

## Results

### FXR activity is decreased in adenine and Alport mice

We first authenticated two FXR antibodies, key biological resources used in this study (**Supplemental Figure 1**). Renal FXR expression and activity were then quantified in models of CKD analogous to APRT deficiency (adenine mice) and Alport syndrome (Alport mice) (**Supplemental Figures 2A and 2B, Supplemental Tables 1 and 2**). Unexpectedly, renal FXR protein expression was unchanged in either model (**Supplemental Figures 2C and 2D**). However, the canonical FXR target gene of Nr0b2 was decreased in both models, thus implying reduced FXR activity (**Supplemental Figures 2E and 2F**).

### FXR agonism reduces CKD in male and female adenine mice

Because renal FXR activity is decreased in the adenine model of CKD, we hypothesized that FXR agonism would reduce disease severity (**Figure 1A**). Adenine mice rapidly lost weight compared to control mice, despite no overt and consistent difference in food intake (**Supplemental Figure 3, Supplemental Table 3**). Male and female adenine mice had increased blood urea nitrogen (BUN) and plasma creatinine compared to control mice, and OCA reduced BUN in female mice and plasma creatinine in both sexes (**Figure 1B**).

**Figure 1:**
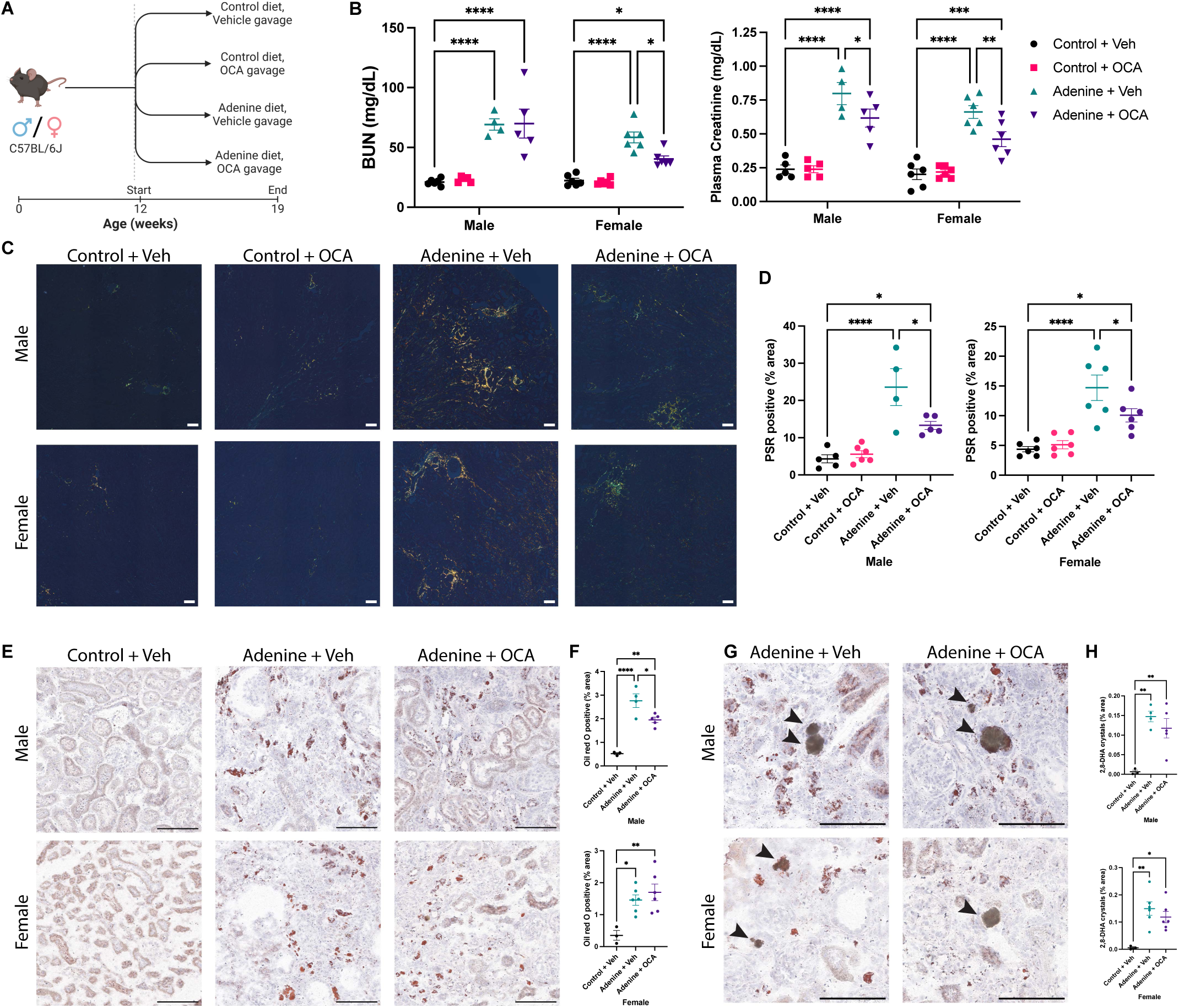
FXR agonism prevents kidney disease in male and female adenine mice. (**A**) Experimental design: OCA treatment was investigated in both sexes in the adenine diet model. (**B**) OCA reduced plasma markers of kidney disease in both sexes. (**C,D**) Representative images and quantification of picrosirius red (PSR) stained kidneys imaged with polarized light show that OCA reduced renal fibrosis in both sexes. (**E,F**) OCA reduced renal lipid accumulation in male but not female mice. (**G,H**) OCA does not affect 2,8-DHA crystal (arrow heads) formation in either sex. Scale bars represent 100 µm. *P < 0.05, ** P < 0.01, *** P < 0.001, **** P < 0.0001. Significance was determined by 2-way (B) and 1-way (**D,G,H**) ANOVA with the Holm-Šídák correction for multiple comparisons. Data are expressed as the mean ± SEM.

Histopathological staining confirmed the treatment success. Adenine mice had increased renal fibrosis compared to control mice, and it was reduced by OCA in both sexes (**Figures 1C, 1D, and Supplemental Figure 4A**). Oil red O staining revealed that adenine mice had increased kidney lipid accumulation in both sexes, and FXR agonism reduced this in male mice only (**Figures 1E, 1F, and Supplemental Figure 4B**). Renal 2,8-dihydroxyadenine (2,8-DHA) crystals were quantified, but there was no difference between vehicle or treatment groups, suggesting that the treatment benefit is not secondary to reduced crystal deposition (**Figures 1G, 1H, and Supplemental Figure 4C**). When taken together, these data provide unequivocal support for the nephroprotective effect of FXR agonism in both sexes in this model of APRT deficiency.

### FXR agonism reduces CKD in Alport mice

Because renal FXR activity is decreased in the Alport model of CKD (**Supplemental Figure 2**), we hypothesized that FXR agonism would reduce disease severity (**Figure 2A**). Within the Col4a3^-/-^ genotype, OCA reduced BUN, plasma creatinine (trend, P < 0.06), and the urinary albumin-to-creatinine ratio (ACR) (**Figure 2B**). OCA also reduced renal fibrosis and renal cortical fibronectin expression (**Figures 2C, 2D, and Supplemental Figure 5A**). In agreement with the albuminuria results, glomerular synaptopodin density was reduced in Alport mice and restored by OCA (**Supplemental Figure 5B**). These data demonstrate the nephroprotective effects of FXR agonism in this model of Alport syndrome.

**Figure 2:**
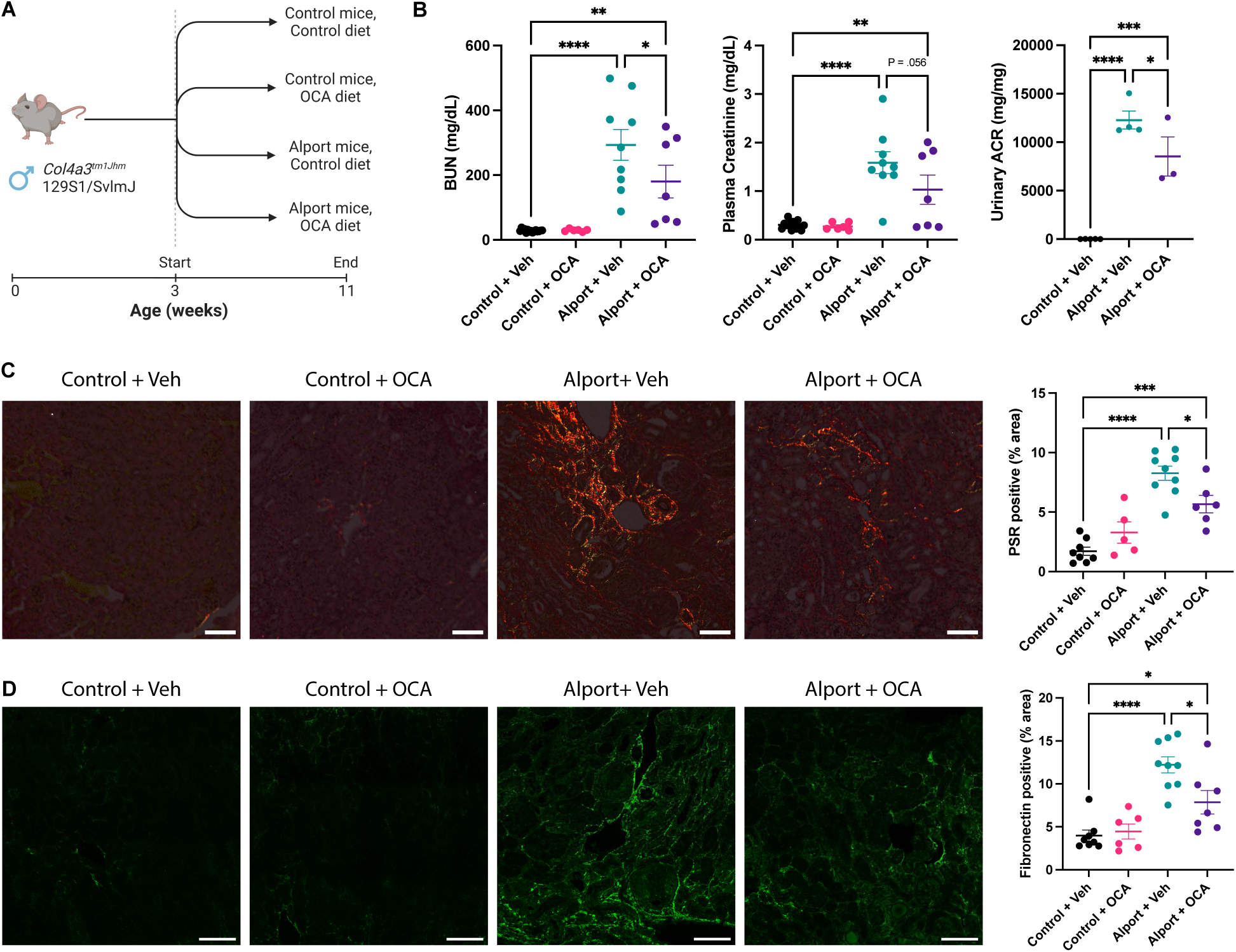
FXR agonism prevents kidney disease in Alport mice. (**A**) Experimental design: Male Alport (Col4a3^-/-^) and control mice were treated with or without OCA. (**B**) OCA treatment reduced blood urea nitrogen (BUN), plasma creatinine (trend), and urinary albumin-to-creatinine ratio (ACR) in Alport mice. (**C**) Representative images and quantification of picrosirius red (PSR) stained kidneys imaged with polarized light show that OCA reduced renal fibrosis. (**D**) Representative images and quantification of fibronectin immunofluorescence further confirm that OCA reduced renal fibrosis. Scale bars represent 100 µm. *P < 0.05, ** P < 0.01, *** P < 0.001, **** P < 0.0001. Significance was determined by 1-way ANOVA with the Holm-Šídák correction for multiple comparisons. Data are expressed as the mean ± SEM.

### FXR agonism reduces renal neutrophilic infiltrate and NETosis in male but not female adenine mice

FXR agonism prevents kidney inflammation, but the mechanistic basis underlying this has not been fully elucidated.^5, 6, 9, 36^ We hypothesized that 2,8-DHA crystals would stimulate NETosis and that FXR agonism would abrogate this via reduced Sphk1 expression.

Adenine mice of both sexes had extensive NETosis, quantified by co-localization of myeloperoxidase (MPO, neutrophil marker) and citrullinated histone H3 (Cit-H3, decondensed DNA) in the absence of CD68 (to rule out macrophage extracellular traps) (**Figures 3A and 3C, arrows**). 2,8-DHA crystals stained non-specifically, and they were excluded during quantitation (**Figure 3A, arrowheads and Supplemental Figure 6**). NETosis would occasionally occur on the periphery of intact 2,8-DHA crystals (**Figure 3C**, arrowheads), and this was included during quantitation. Remarkably, OCA treatment promoted the resolution of neutrophilic inflammation and decreased NETosis in male but not female adenine mice (**Figures 3B and 3D**). The characteristic web-like pattern of colocalized MPO and Cit-H3 can be clearly seen with high magnification confocal microscopy (**Figure 3E**). The arrows highlight areas of extracellular MPO and Cit-H3 colocalization, confirming that this is true NETosis, not just mild citrullination of histone H3 within MPO-positive immune cells. Fluorescence slide scans show the remarkable scale of NETosis in adenine mice (**Figure 3F**).

**Figure 3:**
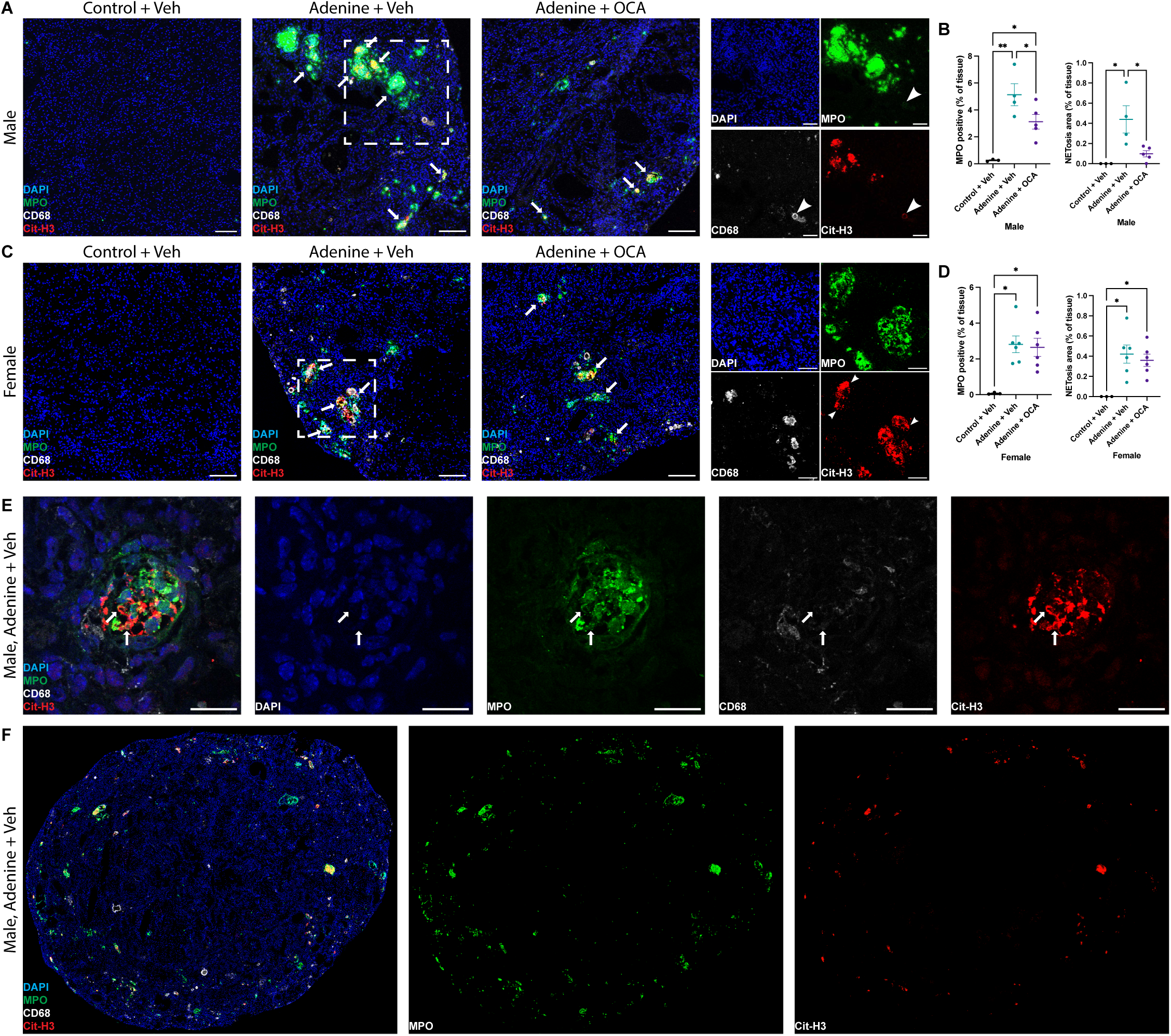
FXR agonism reduces neutrophilic inflammation and NETosis in male adenine mice. (**A**) Male adenine mice have extensive NETosis (arrows) as shown by colocalization (yellow) of myeloperoxidase (MPO, green) and citrullinated histone H3 (Cit-H3, red). (**B**) Quantification of (A) shows that OCA reduced both MPO-positive neutrophilic inflammation and NETosis in male adenine mice. (**C,D**) Female adenine mice have extensive NETosis (arrows) and increased neutrophilic inflammation, but this is unchanged by OCA treatment. 2,8-DHA crystals bind antibodies non-specifically (arrowheads in A inset), and this signal was excluded during quantification. However, NETosis juxtaposed to crystals (arrowheads in C inset) was included in quantification. (**E**) High magnification (63x) confocal microscopy of a male adenine kidney shows characteristic “web-like” NETosis morphology. Arrows in E are aligned pixel-perfectly across channels and represent near-incontrovertible evidence that these lesions are NETs. (**F**) Slide scan (20x) showing the remarkable scale of NETosis in adenine mice. Scale bars represent 100 µm (A,C) and 20 µm (E). *P < 0.05, ** P < 0.01. Significance was determined by 1-way ANOVA with the Holm-Šídák correction for multiple comparisons. Data are expressed as the mean ± SEM.

### FXR agonism reduces renal neutrophilic inflammation and NETosis in male Alport mice

We then investigated male Alport mice to see if FXR agonism also modulated NETosis in that mouse model. Alport mice had extensive neutrophilic infiltrate and NETosis, and both were reduced by FXR agonism (**Figures 4A and 4B**).

**Figure 4:**
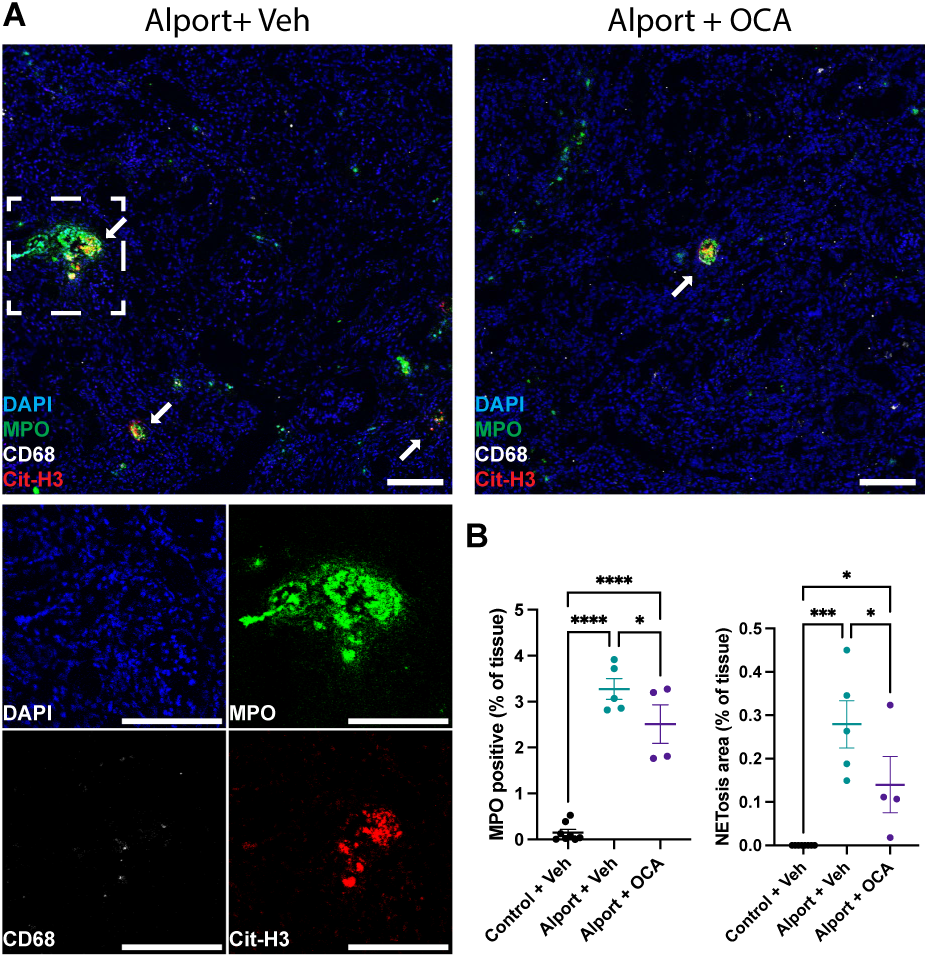
FXR agonism reduces neutrophilic inflammation and NETosis Alport mice. (**A**) Similar to the adenine model, male Alport mice on the nephropathy-susceptible 129S1/SvImJ background have extensive NETosis (arrows) as shown by colocalization (yellow) of myeloperoxidase (MPO, green) and citrullinated histone H3 (Cit-H3, red). (**B**) Quantification of (A) shows that OCA reduced MPO-positive neutrophilic inflammation and NETosis in male Alport mice. Control kidneys were unremarkable (data not shown). Scale bars represent 100 µm. *P < 0.05, *** P < 0.001, **** P < 0.0001. Significance was determined by 1-way ANOVA with the Holm-Šídák correction for multiple comparisons. Data are expressed as the mean ± SEM.

### Neither adenine nor Alport mice have macrophage extracellular traps

Macrophage extracellular traps (METs) have been reported in rhabdomyolysis-induced acute kidney injury.^37, 38^ We did not observe METs in adenine or Alport mice (**Supplemental Figure 7**).

### NETosis occurs in kidney biopsies from humans with Alport syndrome

After confirming the relevance of neutrophils and NETosis in the adenine and Alport models, we then sought to visualize NETosis in human kidney biopsies. NETs were present in 13 of 15 human Alport kidney biopsies, and the degree of NETosis varied between mild, moderate, and severe (**Figure 5, Table 1, and Supplemental Figures 8 and 9**). Kendall rank correlation revealed that NETosis grade was positively correlated with serum creatinine (P < 0.05, τ = 0.67, N = 9). After verifying renal NETosis in patients with Alport syndrome, we then investigated if reduced kidney S1P signaling is a mechanism underlying FXR-mediated inhibition of neutrophilic inflammation and NETosis.

**Figure 5:**
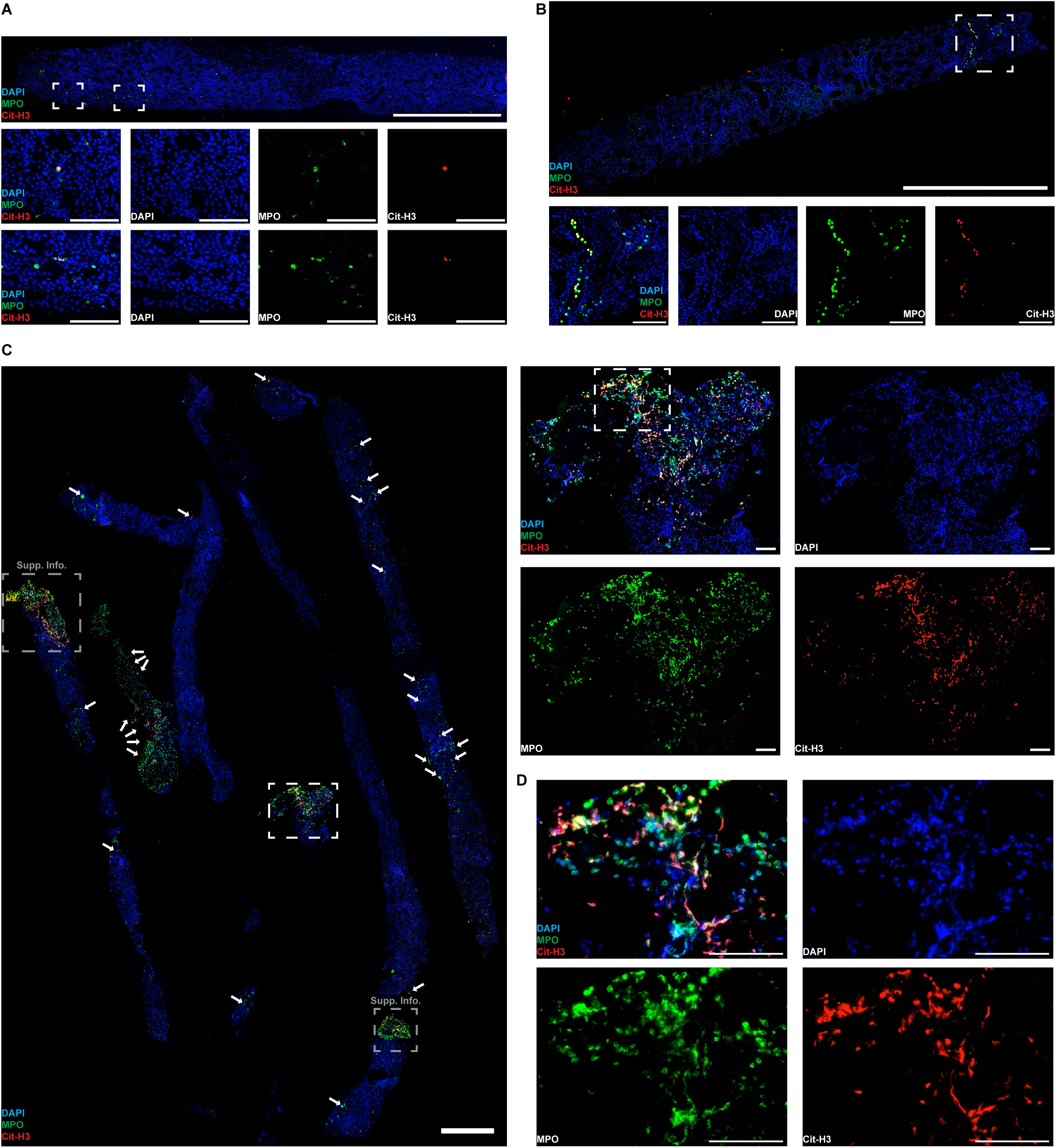
Renal NETosis occurs in humans with Alport syndrome. (**A,B,C**) Kidney biopsies from humans with Alport syndrome were stained for NETosis, identified by colocalization (yellow) of myeloperoxidase (MPO, green) and citrullinated histone H3 (Cit-H3, red). Representative multispectral images (20x) are shown from biopsies with mild (A), moderate (B), and severe (C) NETosis. Only 2 of the 15 kidney biopsies did not have any NETosis (data not shown). Additional inset images from (C) are in the supplemental information. Arrows in (C) indicate areas of significant NETosis that were not also chosen for inset images. Isolated NETting neutrophils are scattered throughout (C) and are unlabeled. (**D**) Multispectral image (40x) showing NETs from inset in (C). Scale bars represent 1 mm in the large images and 100 µm in the inset images. All images in this figure were spectrally unmixed to remove autofluorescence (presented in the supplemental information).

**Table 1:**
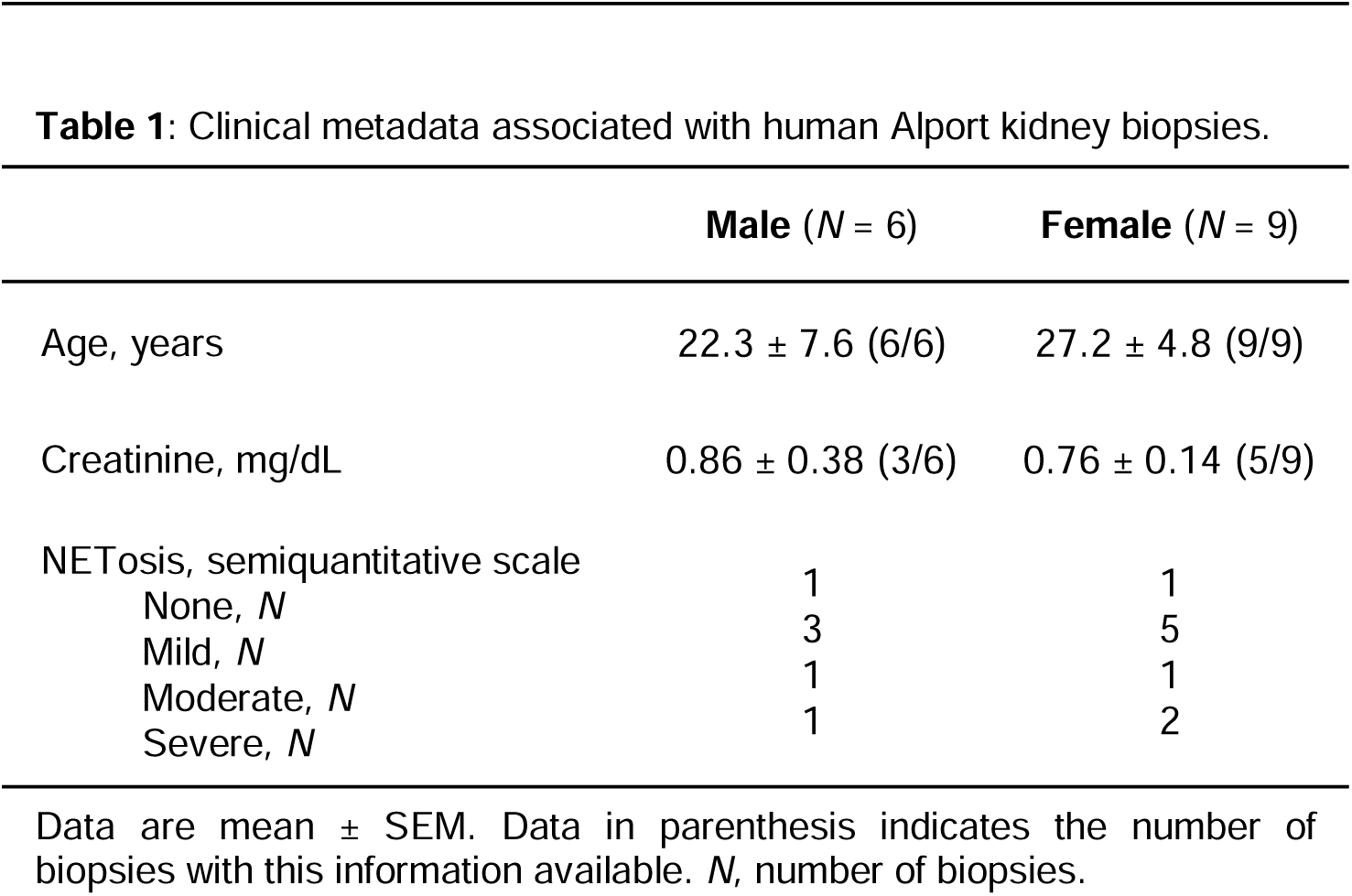
Clinical metadata associated with human Alport kidney biopsies.

|  | **Male** (*N* = 6) | **Female** (*N* = 9) |
| --- | --- | --- |
| Age, years | 22.3 ± 7.6 (6/6) | 27.2 ± 4.8 (9/9) |
| Creatinine, mg/dL | 0.86 ± 0.38 (3/6) | 0.76 ± 0.14 (5/9) |
| NETosis, semiquantitative scale |  |  |
| None, *N* | 1 | 1 |
| Mild, *N* | 3 | 5 |
| Moderate, *N* | 1 | 1 |
| Severe, *N* | 1 | 2 |

Data are mean ± SEM. Data in parenthesis indicates the number of biopsies with this information available. *N*, number of biopsies.

### FXR agonism prevents Sphk1 expression and S1P production in male but not female mice

To determine if FXR modulates inflammation via S1P signaling, we first sought to identify a direct link between reduced FXR activity and renal Sphk1 expression. We quantified Sphk1 in FXR-null mice, and genetic deletion of FXR increased kidney Sphk1 expression (**Figure 6A**). We then quantified Sphk1 expression in our models.

**Figure 6:**
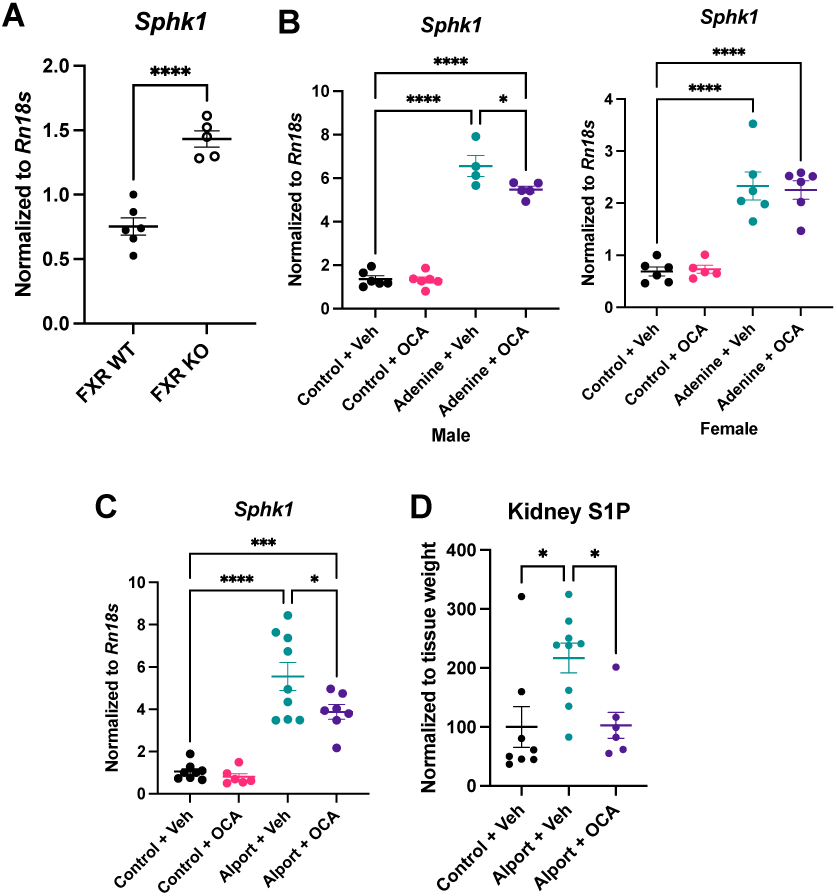
FXR prevents kidney Sphk1 expression and reduces S1P levels in male mice. (**A**) Sphk1 expression is increased upon genetic deletion of FXR in male mice. (**B**) Sphk1 expression is increased in adenine mice of both sexes, and it is reduced by FXR agonism in male but not female mice. (**C**) Similar to that of male adenine mice, Sphk1 expression is increased in male Alport mice and reduced by FXR agonism. (**D**) Analysis of extracted lipids shows that kidney S1P levels are increased in male Alport mice and reduced by FXR agonism. *P < 0.05, *** P < 0.001, **** P < 0.0001. Significance was determined by 1-way ANOVA with the Holm-Šídák correction for multiple comparisons. Data are expressed as the mean ± SEM.

Kidney Sphk1 expression was increased in adenine mice of both sexes and in male Alport mice. Remarkably, FXR agonism decreased Sphk1 expression only in male adenine and Alport mice (**Figures 6B and 6C**). Spearman’s rank correlation revealed a positive association between Sphk1 expression and NETosis in male adenine (trend, P < 0.1) and Alport (significant, P < 0.05) mice, and this was not seen in female mice (**Table 2**). These results are consistent with the sex-dependent prevention of NETosis (**Figures 3 and 4**).

**Table 2:** Spearman correlation between NETosis and Sphk1 expression in adenine and Alport mice.

|  | **Spearman r** | **P value** |
| --- | --- | --- |
| Male adenine (*N* = 9) | 0.600 | 0.097 |
| Female adenine (*N* = 12) | −0.329 | 0.298 |
| Male Alport (*N* = 9) | 0.783 | 0.017 |

*N*, number of mice. Only mice with kidney disease (i.e., adenine or Alport mice) were included in the analysis.

Next, we validated that FXR agonism reduced kidney S1P levels, not just Sphk1 expression. High-performance liquid chromatography tandem mass spectrometry (LC-MS/MS) on extracted kidney lipids revealed that renal S1P in Alport mice was over twice that of control mice, and FXR agonism abrogated this increase (**Figure 6D**).

### Inhibition of de novo sphingosine synthesis reduces renal neutrophilic infiltrate and NETosis

We then sought to establish a causal link between inhibiting S1P signaling and reducing renal neutrophilic infiltrate and NETosis in CKD. To this end, Alport mice with established kidney disease were treated with myriocin to block de novo sphingosine synthesis (**Figure 7A**). There was no difference in kidney disease between Alport groups prior to the start of treatment, and short-term treatment with myriocin did not reduce BUN, plasma creatinine, or albuminuria (Table 3). Nevertheless, there was a remarkable decrease in neutrophils and NETosis in the myriocin-treated mice (**Figure 7B and 7C**). These results show that short-term inhibition of de novo sphingosine synthesis prevents neutrophilic inflammation and NETosis, and that the observed effects in the FXR agonism studies are not just secondary to reduced kidney disease.

**Figure 7:**
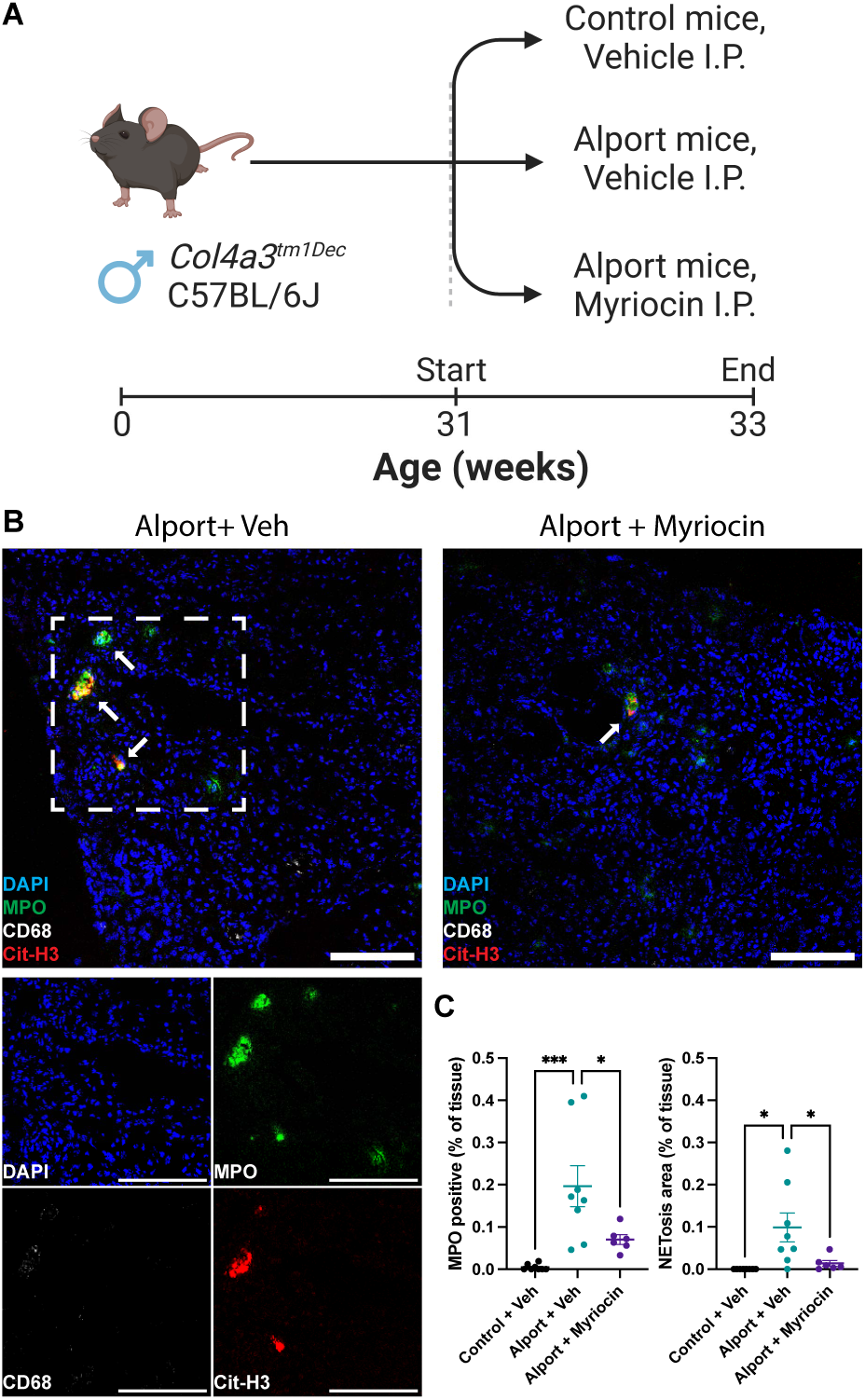
Inhibition of de novo sphingosine synthesis reverses NETosis and promotes resolution of neutrophilic inflammation. (**A**) Experimental design: Male Alport (Col4a3^-/-^) mice on the slow-progressing C57BL/6J background were treated with or without myriocin for two weeks. (**B,C**) Alport mice had moderate NETosis (arrows) which was reversed by short-term inhibition of de novo sphingosine synthesis. Control kidneys were unremarkable (data not shown). Scale bars represent 100 µm. *P < 0.05, *** P < 0.001. Significance was determined by 1-way ANOVA with the Holm-Šídák correction for multiple comparisons. Data are expressed as the mean ± SEM.

**Table 3:** Metabolic parameters for myriocin study in Alport mice.

|  | **Alport + Veh**<br>(*N* = 8) | **Alport + Myr**<br>(*N* = 6) |
| --- | --- | --- |
| **31 weeks of age** |  |  |
| Urinary ACR, mg/mg | 2620 ± 171 | 2590 ± 499 |
| **33 weeks of age** |  |  |
| BUN, mg/dL | 83.7 ± 15.7 | 89.8 ± 25.0 |
| Plasma creatinine, mg/dL | 0.92 ± 0.17 | 1.11 ± 0.34 |
| Urinary ACR, mg/mg | 2374 ± 523 | 2512 ± 374 |

\**P* < 0.05 vs. Alport + Veh. Significance was determined by Student's t-test. Data are expressed as means ± SEM. ACR, albumin-to-creatinine ratio; BUN, blood urea nitrogen; Myr, myriocin; *N*, number of mice; Veh, vehicle.

All together, these results show that FXR agonism represses kidney Sphk1 expression and thus S1P production. In turn, this protects the kidney by reducing both renal neutrophilic inflammation and renal NETosis in a sex-dependent manner (**Figure 8**).

**Figure 8:**
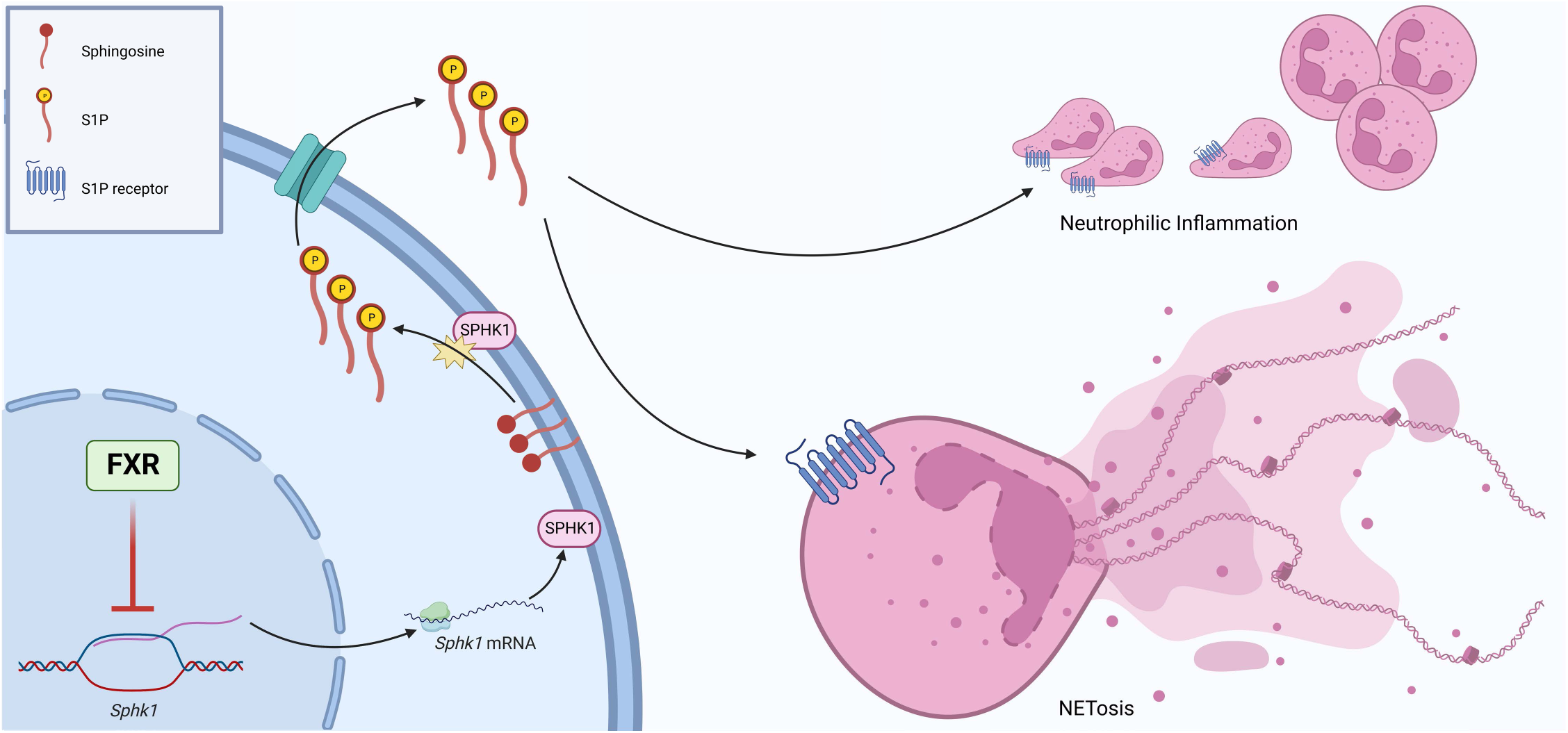
Proposed mechanism for FXR-mediated prevention of neutrophilic inflammation and NETosis in male mice. Under physiological conditions, FXR functions to repress Sphk1 gene expression and maintain kidney S1P at baseline levels. Reduced FXR activity in the setting of chronic kidney disease drives an increase in Sphk1 gene expression, contributing at least in part to elevated renal S1P. This promotes neutrophilic inflammation and NETosis in the kidney. Pharmacological FXR agonism, such as with OCA, represses Sphk1 gene expression and thus prevents these changes in male but not female mice.

## Discussion

We have made several noteworthy advancements presented herein. We are the first to show the benefits of FXR agonism in mouse models of APRT deficiency and Alport syndrome. Additionally, we are the first to show NETosis in these models and in kidney biopsies from patients with Alport syndrome. Finally, we report a novel mechanism whereby FXR agonism reduces renal Sphk1 expression and NETosis in a sex-dependent manner, and inhibition of de novo sphingolipid synthesis prevents NETosis (**Figure 8**).

NETosis is emerging as a critical player involved in the pathogenesis of kidney diseases. There is accumulating evidence for renal NETosis contributing to kidney disease arising from lupus,^39^ hypertension,^40, 41^ vasculitis,^42, 43^ stone formation,^44^ and acute kidney injury.^45^ We are the first to report NETs in mouse models of APRT deficiency and Alport syndrome, as well in kidney biopsies from patients with Alport syndrome.

FXR agonism is known to decrease renal inflammation, but the mechanism underlying this effect is poorly understood. We investigated S1P signaling as a potential mediator of crosstalk between FXR-expressing renal parenchymal cells and neutrophils which do not express detectable levels of FXR on RNA-seq.^46^ In male adenine and Alport mice, Sphk1 expression was increased with kidney disease and reduced with FXR agonism, consistent with recent findings in the liver (supp. info. in ^35^). Furthermore, Sphk1 expression was positively correlated with the severity of NETosis in male but not female mice (Table 2).

Our final key advancement was to establish a causal link between FXR agonism, reduced S1P signaling, and the prevention of neutrophilic inflammation and NETosis. To this end, we blocked de novo sphingosine synthesis in Alport mice with established kidney disease. Myriocin-treated mice had both less neutrophilic infiltrate and fewer NETs than vehicle-treated mice, showing that inhibition of de novo sphingosine synthesis prevents NETosis and promotes resolution of neutrophilic inflammation. Importantly, the short-term treatment with myriocin did not reduce the severity of kidney disease, indicating that this is a direct biochemical mechanism and not solely secondary to the prevention of kidney disease. Our results are consistent with recent reports where inhibition of S1P signaling reduced NETosis and promoted resolution of neutrophilic inflammation in the liver.^23, 25, 27^

In conclusion, FXR agonism protects the kidney in mouse models of APRT deficiency and Alport syndrome. We are the first to report NETs in these mouse models, as well as kidney biopsies from patients with Alport syndrome. Mechanistically, FXR agonism indirectly regulates renal inflammation in a sex-dependent manner by repressing Sphk1 expression in male mice. This inhibits renal S1P signaling, thereby reducing NETosis and promoting resolution of neutrophilic inflammation (**Figure 8**).

## Experimental procedures

### Animal models

Animal studies were approved by the Institutional Animal Care and Use Committee of Georgetown University. All mice were maintained on a 12:12-hr light-dark cycle and fed a grain-based chow (Cat. No. 5053, LabDiet) unless otherwise specified. C57BL/6J mice were obtained from The Jackson Laboratory (Bar Harbor, ME). Col4a3^tm1Jhm^ mice on the 129S1/SvImJ background were obtained as a gift from Dr. Jeffrey H. Miner (Washington University in St. Louis).^47^ Col4a3^-/-^ was the disease genotype, and both Col4a3^+/+^ and Col4a3^+/-^ were the control genotypes. ^tm1Dec^ mice on the C57BL/6J background were obtained as a gift from Sanofi (Framingham, MA).^48, 49^ Genotyping was performed by Transnetyx (Cordova, TN) using RT-qPCR. For the FXR-null experiment, male ^tm1Gonz^ (FXR^-/-^) mice and age matched male C57BL/6 mice were used.^50^ For all animal studies, heparinized plasma and organs were collected upon sacrifice.

### FXR expression and function study in adenine mice

Eight male C57BL/6J mice were purchased at 12 weeks of age, and equal numbers were fed chow alone or admixed with adenine (0.2% w/w) for 7 weeks. Mice were sacrifice at 19 weeks of age.

### FXR expression and function study in Alport mice

Male Col4a3^-/-^ (129S1/SvImJ) and control mice were sacrificed at 8 weeks of age. Urine was collected the week of sacrifice.

### FXR agonism study in adenine mice

Twenty-four male and twenty-four female C57BL/6J mice were purchased at 10 weeks of age and maintained on a purified control diet (D19120401i, Research Diets). At 12 weeks of age, 12 mice of each sex were switched to the control diet admixed with adenine (0.2% w/w, D19120402i, Research Diets), referred to as the adenine diet. Also starting at 12 weeks of age, mice were treated with either vehicle (98% corn oil, 2% DMSO) or OCA (10 mg/kg BW) by oral gavage 5 days per week for 7 weeks. Body weights and food weights were recorded semiweekly. Food intake was calculated as the difference in weight between consecutive datapoints divided by the number of mice in the cage. Each cage represents one datum of food intake. Mice were sacrificed at 19 weeks of age.

### FXR agonism study in Alport mice

Male Col4a3^-/-^ (129S1/SvImJ) and littermate controls were fed chow alone or admixed with OCA (30 mg/kg BW) from 3 weeks to 11 weeks of age. Urine (if present) was collected by directly from the bladder upon sacrifice at 11 weeks of age. Most Alport mice did not have urine to collect.

### Inhibition of de novo sphingosine synthesis in Alport mice

Male Col4a3^-/-^ (C57BL/6J) and heterozygous controls were used. Starting 31 at weeks of age, littermate-matched Col4a3^-/-^ were treated with vehicle (1% DMSO in PBS) or myriocin (1 mg/kg bw) by intraperitoneal injection 6 days per week for 2 weeks. Heterozygous controls were injected with vehicle alone. Urine was collected both before starting the treatment and just before sacrifice at 33 weeks of age.

### Plasma and urine chemistries

BUN was determined using the Jung method.^51^ Plasma/urine creatinine (Cat. No. DICT-500 BioAssay Systems) and urine albumin (Cat. No. 1011, Ethos Biosciences) were determined according to the manufacturers’ instructions.

### RNA isolation and RT-qPCR

Kidney cortex was homogenized in a bead mill homogenizer, and total RNA was isolated with spin columns (Cat. No. 74104, Qiagen). cDNA was synthesized from 100 ng total RNA (Cat. No. 4387406, Applied Biosystems). Real time quantitative PCR was performed (Cat. No. 4385610, Applied Biosystems), and target gene transcript expression was normalized to Rn18s using the ΔΔC_T_ method. Primer sequences are in the supplemental information (Supplemental Table 5).

### Tissue homogenization and immunoblots

A mid-transverse kidney piece was homogenized in T-PER (Cat. No. 78510, Thermo Scientific) with protease and phosphatase inhibitors using a Potter-Elvehjem tissue grinder. Samples of equal protein concentration were prepared in reducing and denaturing conditions (Cat. No. 7722, Cell Signaling Technology).

Twenty-five µg of protein from each sample was ran (100 V, 100 min) on polyacrylamide gels (Cat. No. 5671095, Bio-Rad) using tris-glycine SDS running buffer. Protein was transferred (400 mAh, 60 min, 4 °C) in Towbin buffer onto PVDF membrane (Cat. No. 88518, Thermo Scientific). Total protein was quantified with Ponceau S stain. Membranes were washed with 0.1% Tween 20 in tris-buffered saline (TBST) between each of the following steps. Membranes were blocked (1 hour, 20 °C) with 5% non-fat dry milk (NFDM) in TBST. They were then incubated (overnight, 4 °C) with primary antibodies diluted 1:1000 in 5% bovine serum albumin in TBST. Antibody information is in the supplemental information (Supplemental Table 6). The next day, membranes were incubated with the appropriate secondary antibody diluted 1:5000 in 5% NFDM in TBST. Blots were incubated in ECL substrate (Cat. No. 34580, 34096, or A38554, Thermo Scientific) and imaged (Azure Imager c300, Azure Biosystems).

### Histopathology and immunohistochemistry of FFPE mouse kidneys

Tissues were drop-fixed in 10% neutral-buffered formalin for 24 hours at 4 °C and then transferred to 70% ethanol for storage at 4 °C prior to processing and embedding into paraffin. Tissues were thinly sectioned (2-to-3 µm) onto glass slides. PSR staining was performed according to standard histological procedures. Polarized microscopy on PSR-stained adenine and Alport kidneys was performed using an Olympus model IX83 and a Nikon BioPipeline SLIDE, respectively.

Immunohistochemistry for FXR (1:100, 25 °C, 1 hour) was performed following antigen retrieval (pH9, 95 °C, 1 hour). Staining was performed by Histoserv, Inc. (Germantown, MD). Antibody information is in the supplemental information (Supplemental Table 6). Immunofluorescence of mouse kidneys Mouse kidneys were snap-frozen in liquid nitrogen, embedded in O.C.T., and cryosectioned (5 µm) onto glass slides. Sections were fixed in 4% paraformaldehyde (15-min, 25 °C), blocked in 5% normal donkey serum with 0.3% Triton X-100 (1-hr, 25 °C), incubated with their respective primary antibodies (1:200, overnight, 4 °C), incubated with their respective highly cross-absorbed secondary antibodies (1:400, 1-hr, 25 °C), incubated with DAPI (1:2000, 15-min, 25 °C), and then mounted with Prolong Diamond Antifade Mountant (Invitrogen). Sections were washed in PBS (5-min, 25 °C) in between each of the steps. Antibody information is in the supplemental information (Supplemental Table 6). Kidney sections were imaged in their entirety with a Leica SP8 laser scanning confocal microscope equipped with 20x and 63x objectives.

### Immunofluorescence of FFPE human biopsies

Human Alport kidney biopsies were obtained from the Johns Hopkins University Renal Pathology Archive. Specimens were collected during the course of clinical care and de-identified prior to use in research. Multiplex immunofluorescence for NETosis was performed following antigen retrieval (20-min, pH6, 95 °C). Samples were blocked in 5% normal donkey serum with 0.3% Triton X-100 (1-hr, 25 °C), incubated with their respective primary antibodies (1:100, overnight, 25 °C), incubated with their respective highly cross-absorbed secondary antibodies (1:200, 3-hr, 25 °C), incubated with DAPI (1:2000, 15-min, 25 °C), and then mounted with Prolong Diamond Antifade Mountant (Invitrogen). Sections were washed in PBS (5-min, 25 °C) in between each of the steps. Spectral and autofluorescence controls slides were processed alongside the test slides. Antibody information is in the supplemental information (Supplemental Table 6).

Multispectral microscopy of the human biopsies was performed using a Vectra 3.0 (PerkinElmer) equipped with 20x and 40x objectives. Spectral and autofluorescence control slides were used to construct a spectral library. Using this library, multispectral images were unmixed with inForm (version 2.4.11) into their respective channels (autofluorescence, DAPI, Alexa Fluor 488, and Cy5).

### Tissue lipid extraction and analysis

A transverse kidney piece was homogenized in methanol and extracted overnight in methanol and chloroform at a 2:1 ratio. Samples were then centrifuged, and the supernatant was decanted and evaporated to yield an oily residue of extracted lipids. Lipids were resuspended by sonication in methanol and water at a 1:1 ratio immediately prior to injection.

Relative S1P levels were quantified using high-performance liquid chromatography tandem mass spectrometry as previously described.^52^ Data were analyzed in Analyst (Version 1.6.2), and the relative abundance of S1P was normalized to tissue weight.

### Statistical analyses

Statistical analyses were performed with GraphPad Prism (San Diego, CA) and R programming software. Statistical tests are described in the respective figure captions.

### Disclosures

All authors declare no competing interests.

## Acknowledgements

This research was funded by NIH/NIDDK grants F30DK129003 (BAJ), R01DK116567 (ML), and R01DK127830 (ML); NIH/NCATS grant TL1TR001431 (BAJ); and NIH/NCI grants P30CA016059, P30CA023074, and P30CA051008. The content is solely the responsibility of the authors and does not necessarily represent the official views of the National Institutes of Health. We acknowledge Emma Rowland, Carlos Benitez, and Maria Idalia Cruz for assistance caring for the mice. Experimental design subfigures and **Figure 8** were created with BioRender.com.

