## Supplemental Information for "Farnesoid X receptor agonism prevents neutrophil extracellular traps via reduced sphingosine-1-phosphate in chronic kidney disease"

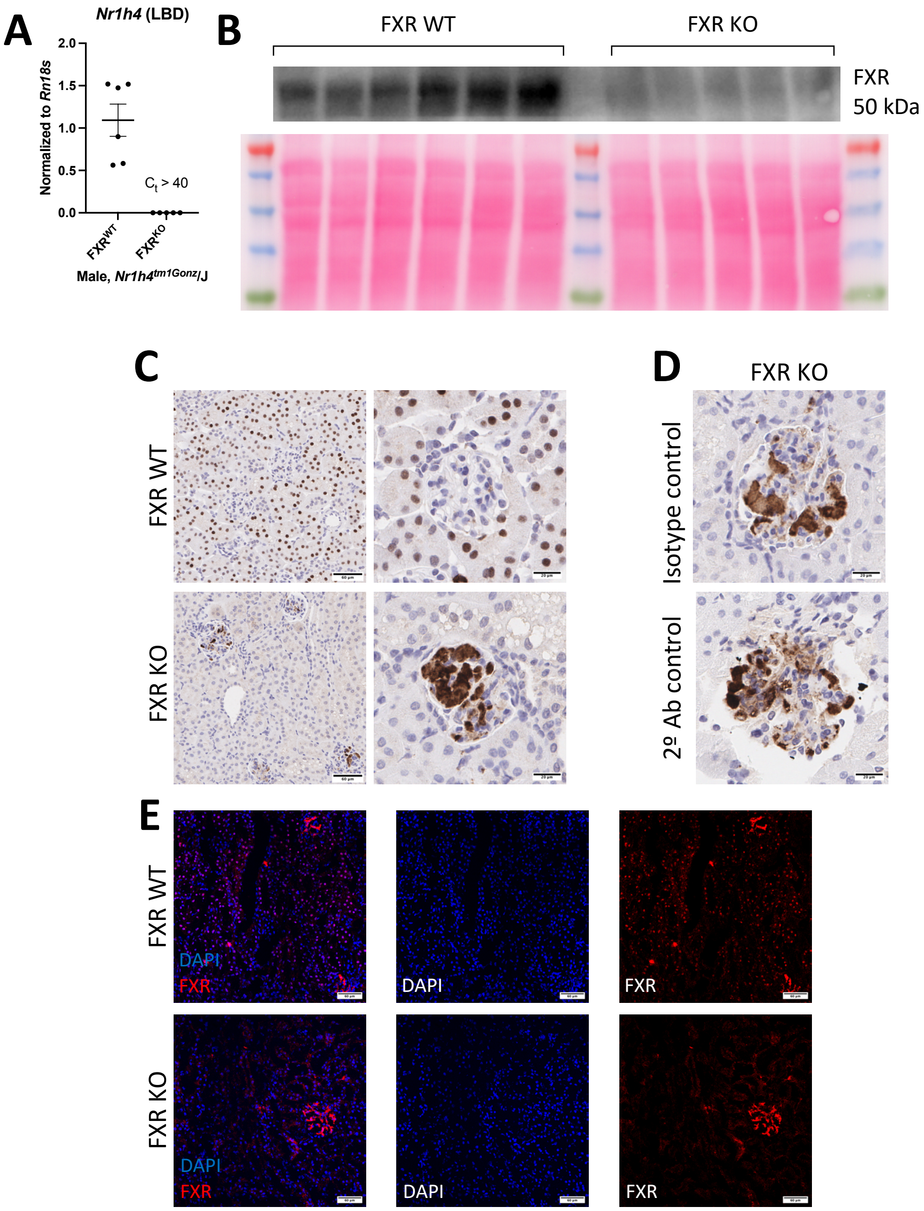

**Supplemental Figure 1: Authentication of key biological resources.** (**A**) Relative transcript level of *Nr1h4* in the kidney using primers specific for the C-terminal ligand-binding domain (LBD) of FXR. $C_t$ values over 40 could not be quantified and are arbitrarily set to zero. (**B**) Immunoblot of total-kidney lysate indicates that FXR is not expressed in $FXR^{-/-}$ mice. (**C**) Immunohistochemistry of control and $FXR^{-/-}$ kidneys shows absence of staining in the tubules of $FXR^{-/-}$ mice. Glomerular signal in $FXR^{-/-}$ kidneys does not exclusively localize to the nucleus. (**D**) Immunohistochemistry of $FXR^{-/-}$ kidneys shows that glomerular signal arises from binding of the secondary antibody to endogenous immunoglobulin. (**E**) Immunofluorescence of shows absence of tubule staining in $FXR^{-/-}$ mice. Scale bars represent 60 µm (tubule images D, E) and 20 µm (glomerular images, D).

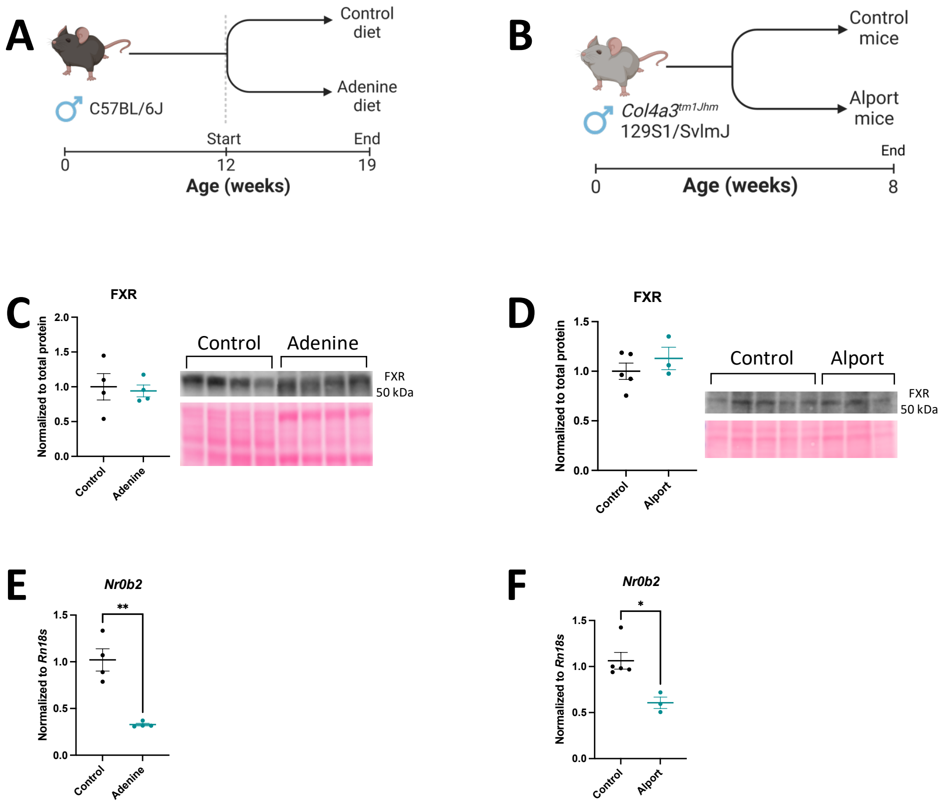

**Supplemental Figure 2: Renal FXR signaling is reduced in adenine and Alport mice.** (**A**) Experimental design: Male mice were fed a control diet or an adenine diet to induce kidney disease. (**B**) Experimental design: Male Alport (Col4a3$^{-/-}$) mice on the fast-progressing 129S1/SvImJ background rapidly develop kidney disease compared to littermate controls. (**C,D**) Renal FXR protein expression was unchanged on immunoblot in adenine (C) or Alport (D) mice compared to healthy controls. Total protein (Ponceau S) was used as a loading control. (**E,F**) Renal transcription of the canonical FXR target gene *Nr0b2* is reduced in adenine (E) and Alport (F) mice, thus implying reduced FXR activity. *P < 0.05, ** P < 0.01. Significance was determined by Student's 2-tailed t test, and data are expressed as the mean ± SEM.

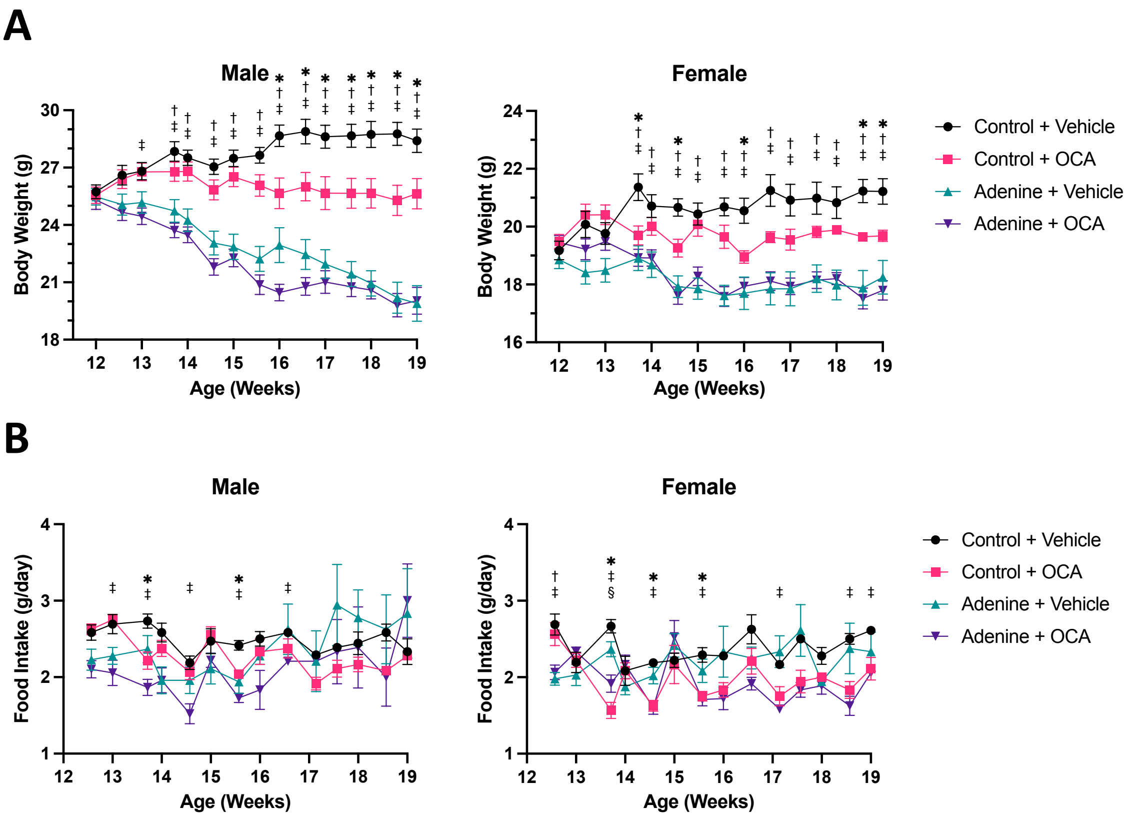

**Supplemental Figure 3: Body weight and food intakes from the FXR agonism study in adenine mice.** (**A**) Body weights during the experiment are plotted over time. (**B**) Average food intakes during the experiment are plotted. The comparisons are as follows: *P < 0.05, control + vehicle vs. control + OCA; †P < 0.05, control + vehicle vs. adenine + vehicle; ‡P < 0.05, control + vehicle vs. adenine + OCA. Significance was determined by a mixed effects model and Holm-Šídák correction for multiple comparisons. Data are expressed as the mean ± SEM.

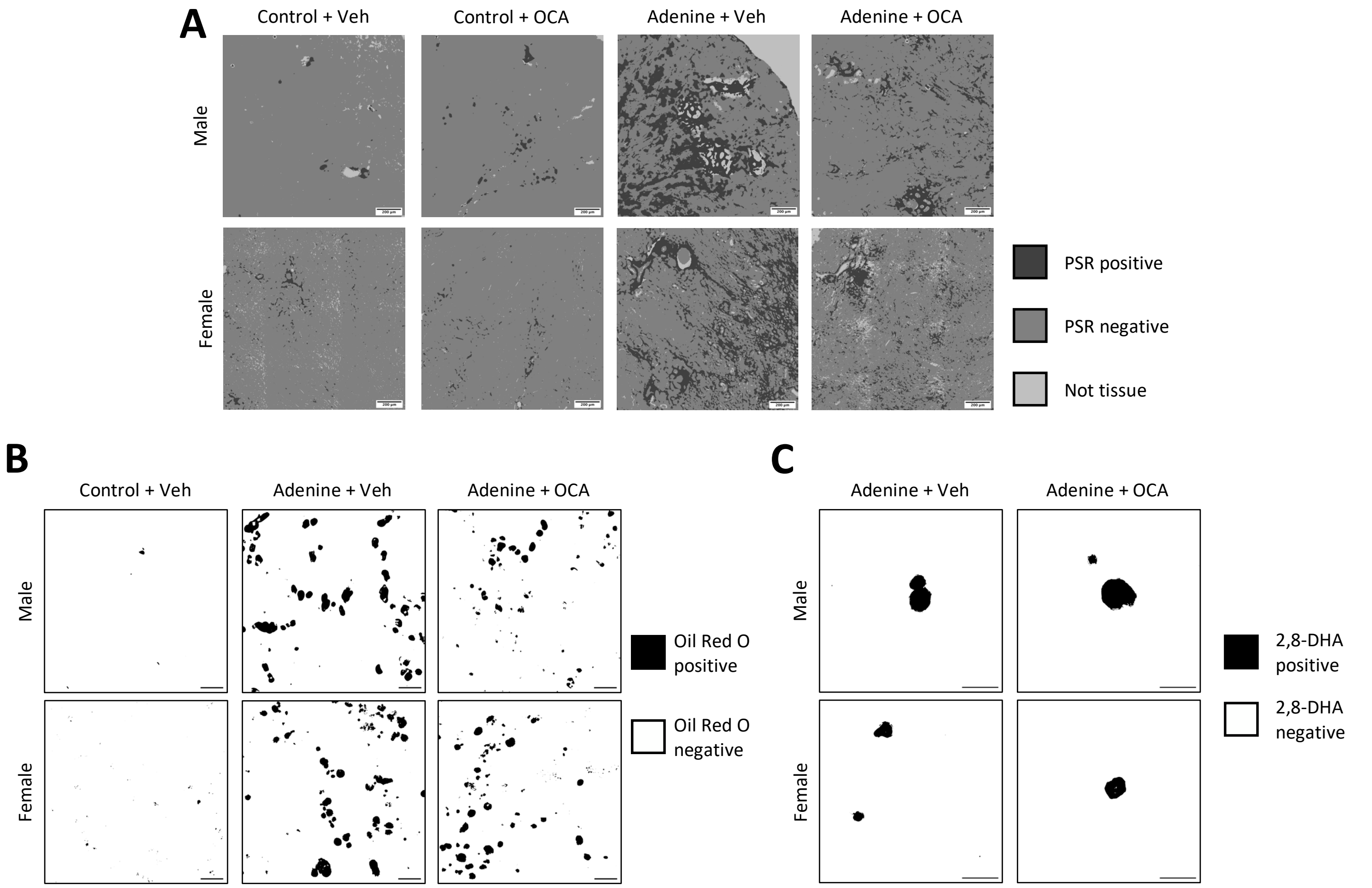

**Supplemental Figure 4: Segmentation of images from the FXR agonism study in adenine mice.** Machine learning was used to identify PSR positive fibrosis, Oil Red O positive neutral lipids, and 2,8-DHA crystals. Segmentation data is shown for the representative images presented in the main text. (**A**) Segmentation of PSR positive tissue from PSR negative tissue and background. Scale bars represent 200 µm. (**B**) Segmentation of Oil Red O positive tissue from Oil Red O negative tissue. Scale bars represent 50 µm. (**C**) Segmentation of 2,8-DHA crystals from surrounding tissue. Scale bars represent 50 µm.

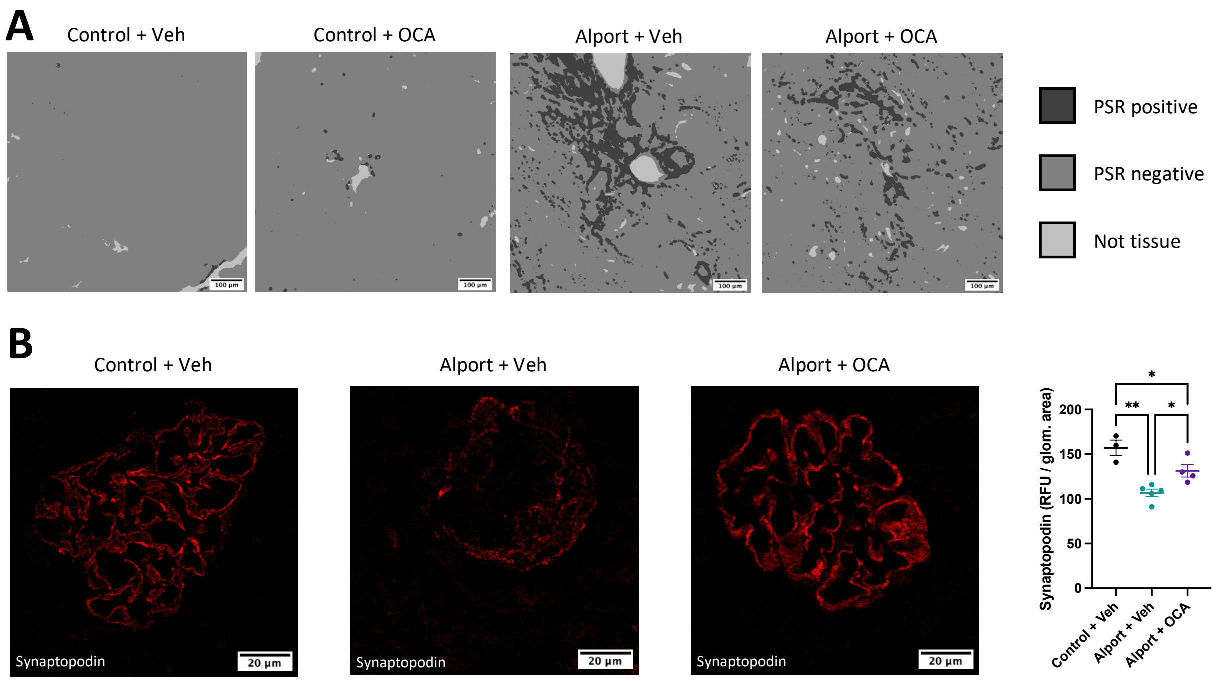

**Supplemental Figure 5: FXR agonism reduces fibrosis and restores glomerular synaptopodin density in Alport mice.** (**A**) Machine learning was used to segment PSR positive fibrosis, PSR negative tissue, and background. Segmentation data is shown for the representative images presented in the main text. Scale bars represent 100 µm. (**B**) Representative images and quantification of synaptopodin immunofluorescence showing that glomerular synaptopodin density was reduced in Alport mice and increased with OCA treatment. Scale bars represent 20 µm. *$P < 0.05$, ** $P < 0.01$. Significance was determined by 1-way ANOVA with the Holm-Šídák correction for multiple comparisons. Data are expressed as the mean ± SEM.

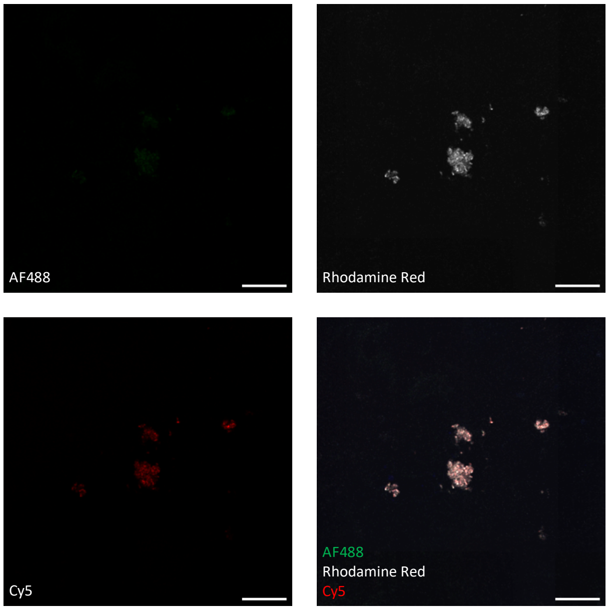

**Supplemental Figure 6: Non-specific fluorescence signal arises from 2,8-DHA crystals.** A secondary-only control section from a male adenine kidney shows that 2,8-DHA crystals contribute to non-specific signal. We were not able to identify blocking conditions that eliminated this signal, and it was manually excluded during quantification of NETs. Scale bars represent 50 µm. AF, Alexa Fluor.

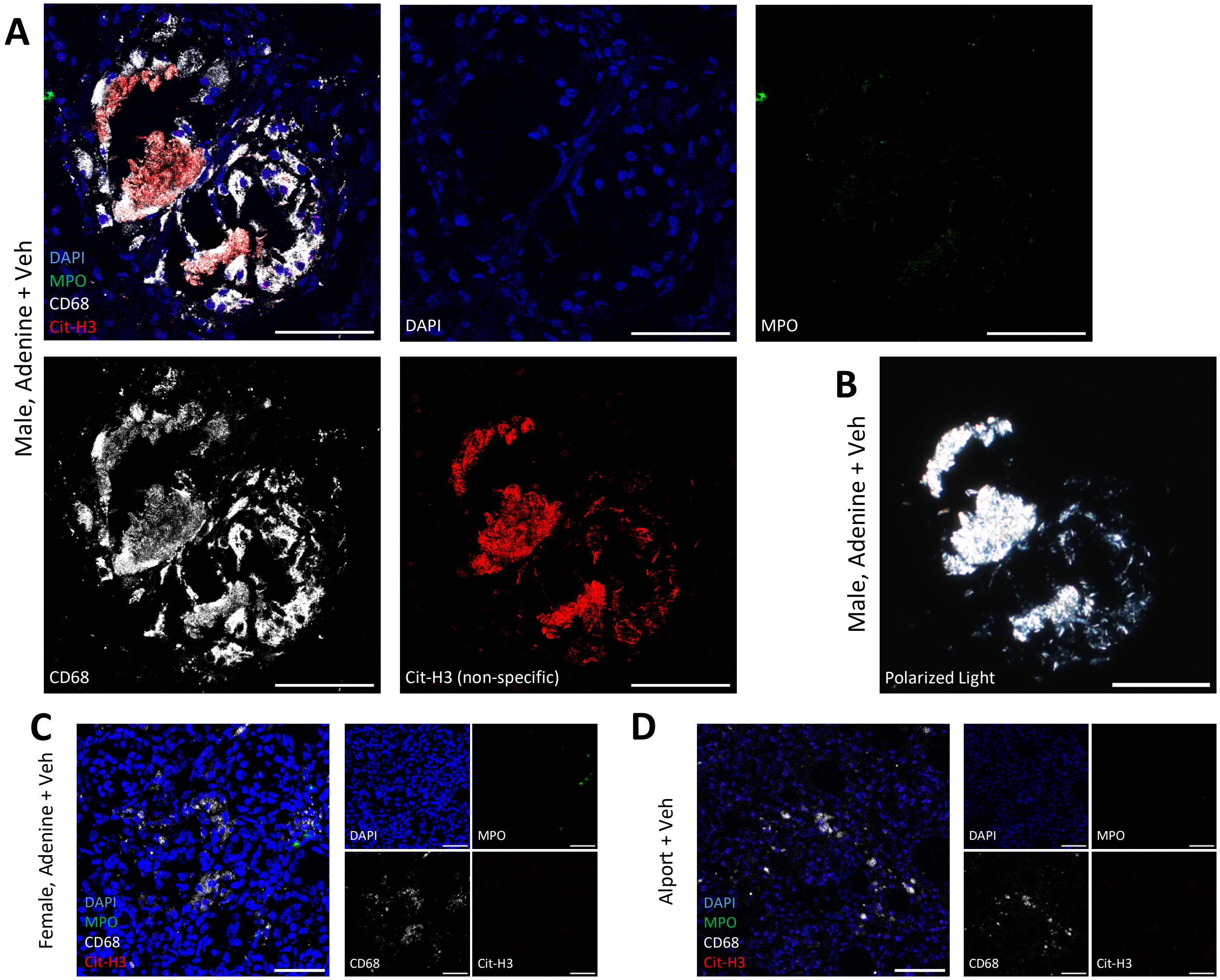

**Supplemental Figure 7: Adenine and Alport mice do not have macrophage extracellular traps.** (**A**) CD68-positive macrophages are present in male adenine mice, but they do not colocalize with specific signal for citrullinated histone H3. Macrophages can be adjacent to large crystals without undergoing METosis (analogous to NETosis, but by macrophages). Macrophages can also phagocytose crystalline shards without undergoing METosis. (**B**) Polarized microscopy confirms that Cy-5 signal (Cit-H3) is localized to the crystal and shards and is thus non-specific signal. (**C**) Female adenine mice also have macrophage clusters that do not undergo METosis. (**D**) Alport mice had macrophage clusters that did not undergo METosis. Control mice from both models did not have macrophage clusters (data not shown). Scale bars represent 50 µm.

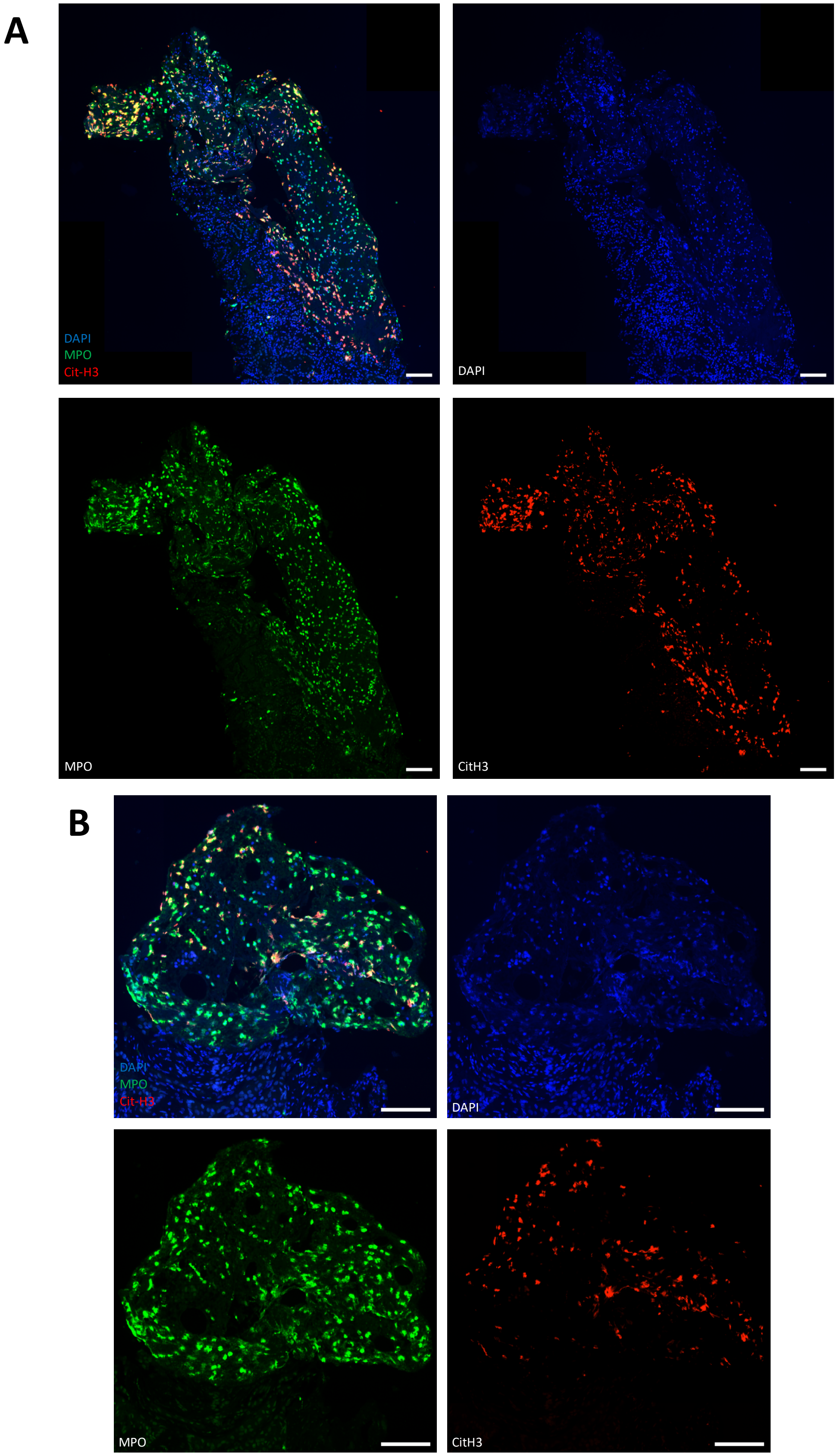

**Supplemental Figure 8: Severe NETosis in a human Alport kidney biopsy.** (**A,B**) Additional inset images of the figure in the main text are shown. Scale bars are 100 um.

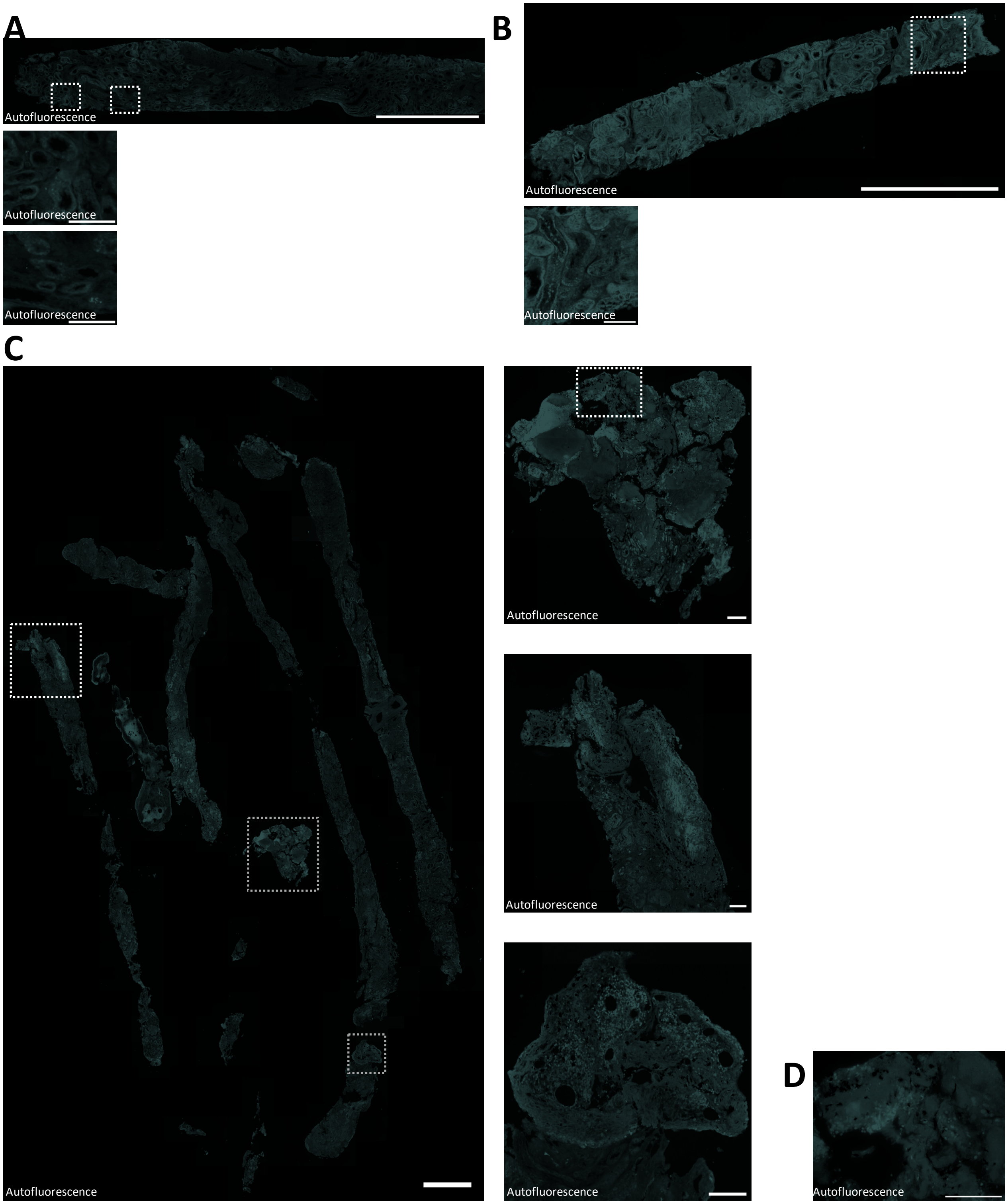

**Supplemental Figure 9: Autofluorescence removed from multispectral images of human Alport biopsies.** (**A,B,C**) Multispectral imaging (20x) was performed on human Alport biopsies stained for NETosis. Autofluorescence that was spectrally unmixed from the representative mild (A), moderate (B) and severe (C) NETosis biopsies are shown. (**D**) Autofluorescence removed from a multispectral image (40x) of the region outlined in the inset image in (C). Scale bars represent 1 mm in the large images and 100 µm in the inset images.

**Supplemental Table 1**: Metabolic parameters at time of euthanasia from the FXR expression and function study in adenine mice.

| | **Control** | **Adenine** |
|---|---|---|
| Body wt, g | 32.8 ± 0.6 | 23.2 ± 0.1 **** |
| Kidney wt / body wt ratio, % | 1.21 ± 0.03 | 1.27 ± 0.02 |
| BUN, mg/dL | 24.0 ± 1.4 | 110.9 ± 5.2 **** |
| Plasma creatinine, mg/dL | 0.23 ± 0.03 | 0.65 ± 0.06 *** |

***$P < 0.001$, ****$P < 0.0001$ vs. control. Significance was determined by Student's t-test. Data are expressed as mean ± SEM ($N = 4$ mice per group). BUN, blood urea nitrogen; wt, weight.

**Supplemental Table 2**: Metabolic parameters at the time of euthanasia from the FXR expression and function study in Alport mice.

| | **Control** (*N* = 5) | **Alport** (*N* = 3) |
|---|---|---|
| Body wt, g | 17.9 ± 0.6 | 18.0 ± 0.5 |
| Kidney wt / body wt ratio, % | 1.39 ± 0.07 | 1.50 ± 0.05 |
| BUN, mg/dL | 29.7 ± 3.1 | 57.6 ± 18.9 |
| Plasma creatinine, mg/dL | 0.18 ± 0.03 | 0.22 ± 0.07 |
| Urinary ACR, mg/mg | 28.2 ± 1.5 | 12127 ± 2975 ** |

**$P < 0.01$ vs. control. Significance was determined by Student's t-test. Data are expressed as mean ± SEM. ACR, albumin-to-creatinine ratio; BUN, blood urea nitrogen; *N*, number of mice; wt, weight.

**Supplemental Table 3**: Metabolic parameters at the time of euthanasia from the FXR agonism study in adenine mice.

| | Control + Veh | Control + OCA | Adenine + Veh | Adenine + OCA |
|---|---|---|---|---|
| **Male** | *N* = 6 | *N* = 6 | *N* = 4 | *N* = 5 |
| Body wt, g | 28.4 ± 0.6 | 25.6 ± 0.8 * | 19.9 ± 0.9 * | 20.0 ± 0.7 * |
| Kidney wt / body wt ratio, % | 1.18 ± 0.08 | 1.20 ± 0.03 | 1.30 ± 0.02 | 1.17 ± 0.04 |
| Hematocrit, % | 51.2 ± 0.6 | 49.1 ± 0.8 | 34.53 ± 1.3 * | 32.5 ± 1.8 * |
| **Female** | *N* = 6 | *N* = 6 | *N* = 6 | *N* = 6 |
| Body wt, g | 21.2 ± 0.4 | 19.7 ± 0.2 * | 18.3 ± 0.6 * | 17.8 ± 0.3 * |
| Kidney wt / body wt ratio, % | 1.15 ± 0.04 | 1.12 ± 0.02 | 0.84 ± 0.05 * | 0.98 ± 0.04 *, # |
| Hematocrit, % | 49.7 ± 0.8 | 48.1 ± 1.1 | 38.9 ± 0.9 * | 41.9 ± 0.6 *, # |

*$P < 0.05$ vs. control + veh. #$P < 0.05$ vs. adenine + veh. Significance was determined by 1-way ANOVA with the Holm-Šídák correction for multiple comparisons. Data are expressed as the mean ± SEM. *N*, number of surviving mice; OCA, obeticholic acid; Veh, vehicle; wt, weight.

**Supplemental Table 4**: Metabolic parameters at the time of euthanasia from the FXR agonism study in Alport mice.

| | **Control + Veh** (*N* = 8) | **Control + OCA** (*N* = 6) | **Alport + Veh** (*N* = 9) | **Alport + OCA** (*N* = 7) |
|---|---|---|---|---|
| Body wt, g | 22.8 ± 1.1 | 22.1 ± 0.9 | 15.1 ± 0.5 **** | 16.5 ± 0.7 **** |
| Kidney wt / body wt ratio, % | 1.29 ± 0.02 | 1.41 ± 0.06 | 1.34 ± 0.04 ### | 1.64 ± 0.06 **** |

****$P < 0.0001$ vs. control + veh. ###P < 0.001 vs. Alport + veh. Significance was determined by 1-way ANOVA with the Holm-Šídák correction for multiple comparisons. Data are expressed as the mean ± SEM. *N*, number of surviving mice; OCA, obeticholic acid; Veh, vehicle; wt, weight.

**Supplemental Table 5:** Nucleotide sequences for primers.

| | Strand | Sequence |
|---|---|---|
| *Nr0b2* | Forward | CTACCCTCAAGAACATTCCAG |
| | Reverse | GGCACCAGACTCCATTCC |
| *Nr1h4* (LBD) | Forward | CACGCTGAGATGCTGATGTC |
| | Reverse | ACCAGGTACAAACGAAACAAGG |
| *Rn18s* | Forward | CGGCTTAATTTGACTCAACAC |
| | Reverse | ATCAATCTGTCAATCCTGTCC |
| *Sphk1* | Forward | GCCACCTCCAGAAGAACC |
| | Reverse | ACTTTAGAAATAACCTCCCATA |

LBD, ligand binding domain.

**Supplemental Table 6**: Antibody supplier and application information.

| Target | Host | Conjugate | Supplier | Catalog number | Application |
|---|---|---|---|---|---|
| Citrullinated histone H3 | Rb | – | Abcam | ab5103 | IF |
| CD68 | Rt | – | Bio-Rad | MCA1957 | IF |
| Fibronectin | Rb | – | Sigma | F3648 | IF |
| FXR | Ms | – | Cell Signaling | 72105 | IB |
| FXR | Ms | – | R&D Systems | PP-A9033A-00 | IHC, IF |
| MPO | Gt | – | R&D Systems | AF3667 | IF |
| Synaptopodin | Rb | – | Sigma | S9442 | IF |
| Gt IgG | Dk | AF 488 | Invitrogen | A11055 | IF |
| Ms IgG | Gt | AF 594 | Invitrogen | A11032 | IF |
| Ms IgG, light chain specific | Gt | HRP | Millipore | AP200P | IB |
| Ms IgG, native chain | Rt | HRP | Rockland | 18-8817-30 | IB |
| Rb IgG | Dk | Cy5 | Jackson Immunoresearch | 711-175-152 | IF |
| Rt IgG | Dk | Rhodamine red | Jackson Immunoresearch | 712-295-153 | IF |

AF, Alexa Fluor; Dk, donkey; Gt, goat; HRP, horseradish peroxidase; IB, immunoblot; IF, immunofluorescence; IHC, immunohistochemistry; Ms, mouse; Rb, rabbit; Rt, rat.
